# H3K27me3 is dispensable for early differentiation but required to maintain differentiated cell identity

**DOI:** 10.1101/2020.06.27.175612

**Authors:** Sara A. Miller, Manashree Damle, Robert E. Kingston

## Abstract

Polycomb repressive complex 2 (PRC2) catalyzes trimethylation of histone H3 on lysine 27 and is required for normal development of complex eukaryotes. The requirement for H3K27me3 in various aspects of mammalian differentiation is not clear. Though associated with repressed genes, the modification is not sufficient to induce gene repression, and in some instances is not required. To examine the role of the modification in mammalian differentiation, we blocked trimethylation of H3K27 with both a small molecule inhibitor, GSK343, and by introducing a point mutation into EZH2, the catalytic subunit of PRC2. We found that cells with substantively decreased H3K27 tri-methylation were able to differentiate, which contrasts with EZH2 null cells. Different PRC2 targets had varied requirements for H3K27me3 in repressive regulation with a subset that maintained normal levels of repression in the absence of methylation. The primary cellular phenotype when H3K27 tri-methylation was blocked was an inability of the altered cells to maintain a differentiated state when challenged. This phenotype was determined by H3K27me3 deposition both in embryonic stem cells and in the first four days of differentiation. H3K27 tri-methylation therefore was not necessary for formation of differentiated cell states but was required to maintain a stable differentiated state.

## Introduction

Polycomb repressive complex 2 (PRC2) is a highly conserved protein complex that is required for proper axial patterning of vertebrates. It is comprised of the core subunits EZH2, SUZ12, EED and RBAP48, with additional subunits some of which are cell-type or developmental stage specific (Healy et al., 2019; Margueron & Reinberg, 2011; Shen et al., 2009; van Mierlo et al., 2019). The complex is crucial for differentiation, but it is not required for the self-renewing phenotype associated with embryonic stem (ES) cells (Aloia et al., 2013; Chamberlain et al., 2008). At a molecular level, PRC2 family complexes are the only histone 3 lysine 27 (H3K27) methyltransferases identified in mammals. EZH2 is the primary catalytic component of these complexes, with its paralog EZH1 also contributing to catalytic activity in some instances (Margueron et al., 2008; Shen et al., 2008; Wassef et al., 2019; Xu et al., 2015). Lysine residues can be mono-di- or tri-methylated, with PRC2 able to catalyze all levels of methylation. Increased methylation is generally associated with gene repression, and H3K27 tri-methylation (H3K27me3) is a key marker of facultative heterochromatin. However, the tri-methylation of H3K27 is neither required nor sufficient to induce gene repression (Ahmed et al., 2018; Chamberlain et al., 2008; Ferguson et al., 2018; O’Geen et al., 2017). Dissecting the role of tri-methylated H3K27 in gene regulation is important for understanding how genes become repressed during development and retain their repression in differentiated cells.

Changes in expression level or activity of Polycomb complexes have developmental and disease effects. Both hyper-activating and inactivating mutations in PRC2 have been identified from a variety of human cancers (Basheer et al., 2019; Cyrus et al., 2019; Jain & Di Croce, 2016). Malignancies with both types of alterations to PRC2 function have been linked to cancer progression and poor prognosis in a wide variety of tissue (Abdel Raouf et al., 2019; Basheer et al., 2019; Bohm et al., 2019; Bremer et al., 2019; Deng et al., 2019; Dou et al., 2019; Karlowee et al., 2019; Krill et al., 2020; Matsubara et al., 2019; Mechaal et al., 2019; Shi et al., 2019; Tian et al., 2019; Wasenang et al., 2019; M. J. Zhang et al., 2019; Q. Zhang et al., 2019). These findings are consistent with the initial identification of Polycomb-Group genes in Drosophila where haploinsufficiency yielded developmental phenotypes (Lewis & Mislove, 1947; Schuettengruber et al., 2017). Inserting these disease mutations into cells alters the modification profiles of H3K27 and can change their developmental potential. Increased methylation can often push cells toward a neuro progenitor phenotype (Juan et al., 2016; Pasini et al., 2007; Thornton et al., 2014). In cancer patients, increased levels of expression of PRC2 components and especially EZH2 leads to hypermethylation and is associated with poor prognosis (Tian et al., 2019; Wu et al., 2019). Consequently, there is significant interest in targeting this complex pharmaceutically. Small molecules that inhibit PRC2 methyltransferase activity have been approved for clinical trials (Fioravanti et al., 2018; Kondo, 2014; Liu et al., 2015; Lue & Amengual, 2018; Qi et al., 2012; Shi et al., 2019; Xu et al., 2015; Yamagishi & Uchimaru, 2017; Yang et al., 2019). It remains, however, unclear how far this activity can be manipulated before risking adverse effects from having too little of the modification present. The ambiguity of the role of H3K27me3 in disease progression increases the importance of understanding the mechanisms by which PRC2 methylation activities regulate gene expression during development.

Previous work on PRC2 has shown that the complex is dispensable for the propagation of self-renewing, undifferentiated cells that are phenotypically indistinguishable from WT ES cells. These studies have also shown that PRC2 deficient cells do not differentiate properly (Chamberlain et al., 2008; Lavarone et al., 2019; Pasini et al., 2007; Pasini et al., 2004; Shen et al., 2008). Since the entire complex is disrupted when any of the core components are knocked out, it is not clear whether it is solely the lack of methylation that is causing these developmental defects, or whether there are other mechanistic roles PRC2 plays that are distinct from methylation(Collinson et al., 2016; Rai et al., 2013; Shan et al., 2017; Yu et al., 2017). A recent study examined the impact of a mutation in *Ezh2* that blocks all levels of methylation. When these cells were differentiated into embryoid bodies they showed visual phenotypic differences (Lavarone et al., 2019), demonstrating that complete removal of the methyltransferase activity of PRC2 impairs normal differentiation. We focus here on the developmental phenotypes caused by specific loss of the H3K27me3 modification, as opposed to loss of all methylation.

To determine the contribution that H3K27me3 makes to gene repression, we examined cells with a point mutation within the SET domain of *Ezh2* that generates a hypomorph that is predominately defective in tri-methylation. We also analyzed cells treated with a small molecule inhibitor that blocks tri-methylation more efficiently than di-methylation. The expression level of PRC2 has known effects on gene regulation, so we used strategies that do not impact the protein level or complex integrity, thus allowing a focus on methyltransferase activity. There are multiple small molecules that interrupt PRC2 methyltransferase activity both for research and current clinical trials; we used GSK343 (Bradley et al., 2014; Fraineau et al., 2017; Liu et al., 2015; Xu et al., 2019; Yang et al., 2019). Mutations in the catalytic SET domain of *Ezh2* have been identified from a variety of cancers; we created cell lines containing one of the inactivating mutations for our experiments (Antonysamy et al., 2013). We found that this mutation had a strong impact on H3K27me3 levels, but not on H3K27me2 levels. When these cells were differentiated in an undirected fashion they did not show the significant visual phenotypes seen with *Ezh2* knockout cells. While expression of some genes was altered, there were a number of PRC2 target genes that did not rely upon H3K27me3 for their regulation. The most dramatic phenotype observed was when cells were challenged to maintain their differentiated identity. Inhibitor treated and point mutated cells reverted readily to an ES phenotype when placed back into conditions that support ES cell growth. Thus, H3K27me3 was required for cells to maintain their identity rather than for the initial differentiation.

## Results

PRC2 catalyzes di- and tri-methylation of histone H3 and is associated with gene repression. Previous studies have shown that deletion of PRC2 components in mouse ES cells has minimal effects on the undifferentiated, self-renewing state (Juan et al., 2016; Lavarone et al., 2019; Shen et al., 2008; Wassef et al., 2019). Yet deletion of EZH2, the catalytic subunit of PRC2, results in cells that do not differentiate and embryos that are reabsorbed by day E10.5 (Pasini et al., 2004). However, the requirements for specific methylation functions of PRC2 in differentiation and in the accompanying changes in gene expression have not been explored in detail. We examined the role for H3K27me3 in regulation by looking at stochastic differentiation of ES cells into embryoid bodies. Embryoid body (EB) formation is a well-studied method of undirected differentiation with predictable changes in gene expression (Fig 1A and (Behringer et al., 2016; Dang et al., 2002). Cells where PRC2 core components haven been deleted fail to form normal embryoid bodies, presumably because the cells apoptose when pushed to differentiate(Chamberlain et al., 2008). Since PRC2 has been shown to play regulatory roles independent of H3K27 tri-methylation, including H3K27 di-methylation and proposed non-methyltrtansferase functions, the requirement for the H3K27me3 modification at individual genes during differentiation as well as its role in maintaining cell identity remains unclear (Ahmed et al., 2018; Ai et al., 2017; Pereira et al., 2010).

**Figure 1.**
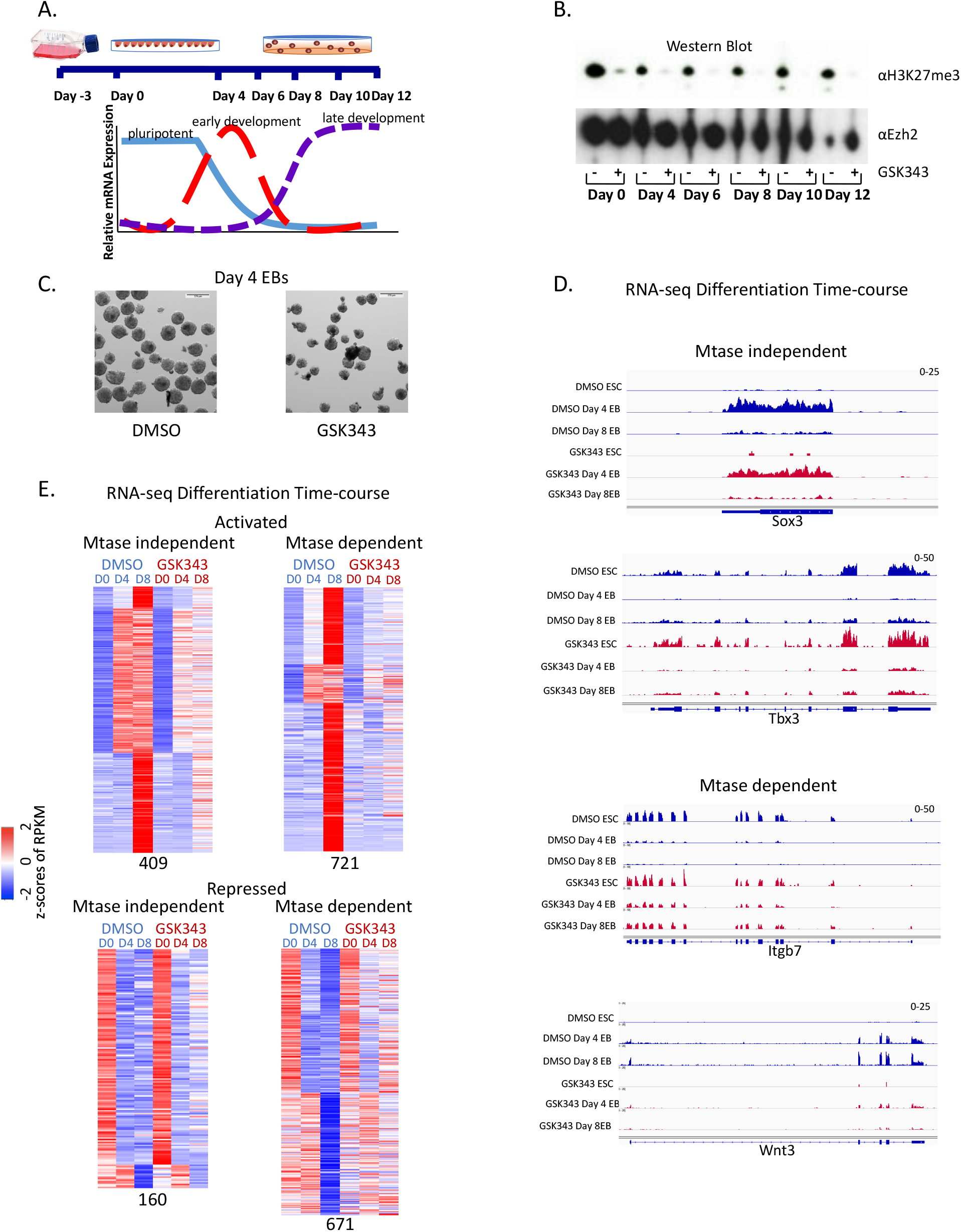
(A) Diagram of embryoid body (EB) formation time-course. (B) Western blot of cells treated with GSK343 EZH2 inhibitor or DMSO control probed with H3K27me3 and EZH2 antibodies. (C) 54x magnification of day 4 EBs from cells treated with either DMSO control or GSK343. (D) Screen shots from RNA-seq over the EB differentiation time-course from Mtase independent (Sox3 and Tbx3) and Mtase dependent (Itgb7 and Wnt3) genes from cells treated with DMSO (blue) or GSK343 (red). (E) Heatmap showing expression patterns of Mtase dependent and independent genes in untreated (DMSO) and treated (GSK343) ES cells over development time-course. Z-scores of average RPKMs of duplicates are shown for each gene across all samples. Genes are separated based on if they were activated (top) or repressed (bottom) in untreated cells over time as well as if they had a similar expression pattern in treated cells (Mtase dependent genes) (left) or a different expression pattern (Mtase independent genes) (right). The four groups are further clustered based on fold-changes over time in untreated cells.

### Inhibiting methyltransferase activity with the small molecule GSK343 does not block differentiation potential

As a starting point to understand the role of H3K27 methylation by PRC2 during ES cell differentiation, we treated cells with the small molecule inhibitor GSK343 which blocks the methyltransferase activity of EZH2 (Fioravanti et al., 2018; Lue & Amengual, 2018; Yang et al., 2019). Treatment with DMSO was used as a control (Fig 1B). Unlike PRC2 knockouts, GSK343 treated cells form embryoid bodies that have a similar morphology to their DMSO treated counterparts (Fig 1C.) Embryoid bodies formed after either DMSO or GSK343 treatment are phenotypically similar to those made by WT cells. In contrast, PRC2 knockout cells tested in parallel failed to form embryoid bodies ((Lavarone et al., 2019) and Fig. 2E). Thus, the inhibition of methyltransferase activity can be phenotypically separated from PRC2 knockout cells.

**Figure 2.**
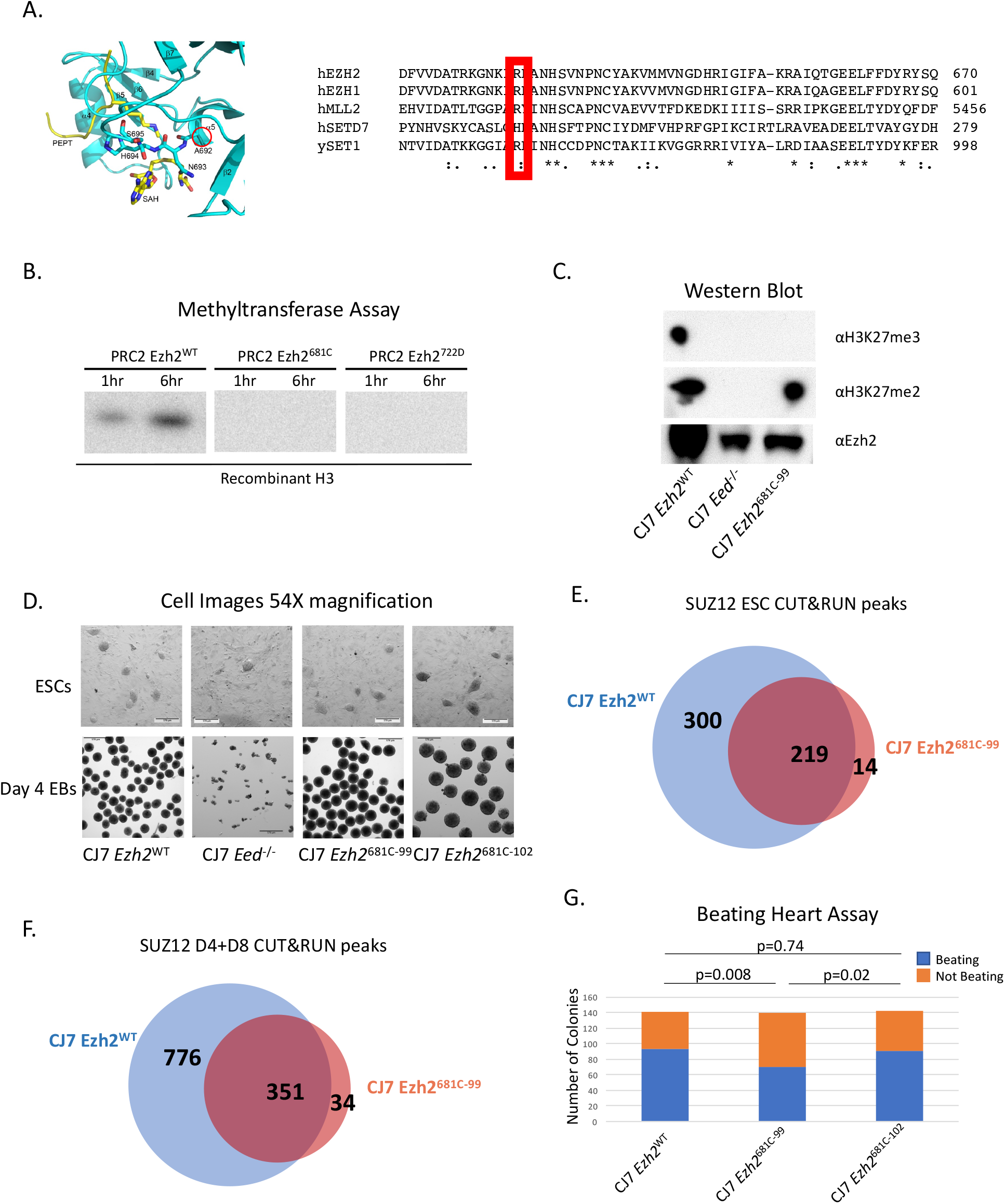
(A) Diagram of *Ezh2* SET domain structure with the 681 residue circled and an alignment of SET domains with the analogous residue highlighted in red adapted from (Antonysamy et al., 2013). It is highly conserved across species and methyltransferases. (B) In vitro methyltransferase assay with PRC2 comprised of WT or mutant EZH2, EED, SUZ12, and RBAP48. Recombinant H3 is the substrate and a radioactive SAM was the methyl donor. Reactions progressed for 1 or 6 hours. (C) Western blot from WT, mutant and PRC2 knockout cells probed with H3K27me and EZH2 antibodies. (D) 54x magnification of embryonic stem cells (ESCs) or embryoid bodies that have differentiated for 4 days (Day 4 EBs) from WT, 681C-99, 681C-102 and *Eed^-/-^* cells. (E) Venn diagram showing the overlap in SUZ12 CUT&RUN peaks from WT and 681C-99 mutant ESCs. (F) Venn diagram showing the overlaps in SUZ12 CUT&RUN peaks from WT and 681C-99 mutant cells differentiated as embryoid bodies for 4 to 8 days. (G) Quantification of beating heart assay from WT, 681C-99 and 681C-102 cells that were differentiated as EBs for 4 days and then individually plated in differentiation media. Blue shows the number of EBs that gave rise to beating cells and Orange shows those that did not.

To determine the molecular phenotype of blocking methyltransferase activity, we performed RNA-seq experiments to compare changes in gene expression between cells treated with the inhibitor and control over the differentiation time course. Gene expression was largely similar between the two treatments especially in early stages of differentiation (Supp. Fig. 1). We then examined known PRC2 target genes and found examples of genes whose expression pattern across differentiation was the same in DMSO and GSK343 treated cells (e.g. Tbx3 or Sox3, Fig. 1D) as well as genes whose expression was altered in GSK343 treated cells (e.g., Wnt3 and Itgb7, Fig 1D). The trends observed with these two classes of genes were found to be mirrored in the full complement of PRC2 targets. We classified PRC2 target genes into two sets, those that required H3K27me3 for their regulation and those that do not need this modification to preserve normal gene expression patterns during embryoid body formation. Genes were defined as independent of the H3K27me3 modification if their expression remained within 1.2 fold of the WT expression at all time points (Fig 1E). We conclude that inhibiting the methylation activity of PRC2 alters gene expression at a subset of targets but does not block differentiation the way knockouts of the complex do.

### Point mutation to the SET domain inhibits methyltransferase activity

We sought to extend and refine the findings made using GSK343 by generating a hypomorphic mutation in *Ezh2* that had a defined impact on H3K27 methylation. We generated mutant cells that allowed us to examine the contribution of H3K27 tri-methylation to gene regulation and differentiation during EB formation. Inactivating point mutations in the SET domain of EZH2 have been identified in several human cancers. We chose one of those mutations, human R635C, which is analogous to mouse R681C, for analysis in mouse cells. This residue is conserved across methyltransferases and located in a region that is important for coordinating the methyl donor (Fig 2A and (Antonysamy et al., 2013). We used a baculoviral system to reconstitute WT and mutant PRC2. We confirmed that this residue is important for tri-methylation of H3K27 by PRC2 using an in vitro methyltransferase assay. We compared it to WT EZH2 and another SET domain mutation (Y722D) that has been previously shown to block methylation (Lavarone et al., 2019), Fig 2B). The complexes that contained WT EZH2 methylated both core histones and purified H3 in a time and concentration dependent manner (Supp. Fig. 2). In contrast, mutant complexes show drastically reduced levels of activity. Using two separate purifications, residual activity of the mutant complex was always under 2% of the purified WT complexes (Fig. 2B and Supp. Fig. 2). This was not due to a failure of the mutant EZH2 to incorporate into a stable complex as all core components were pulled down with a flag-tagged SUZ12 (Supp. Fig. 2E). We conclude that the *Ezh2* 681C mutation reduces the methyltransferase activity of the complex by at least 50-fold.

We introduced the R681C mutation into the CJ7 mouse ES cell line using the CRISPR-Cas9 system. We isolated two independent with this homozygous point mutation in the endogenous *Ezh2* gene (called CJ7 Ezh2^681C-99^ and CJ7 Ezh2^681C-102^). The phenotypes that we describe below were consistent between these two independent clones.

We determined whether the point mutation would lead to significant reduction of H3K27me3 in cells as anticipated from the enzymatic defect seen in vitro. Lysine residues can be mono, di or tri-methylated. Mono-methylation is spread broadly throughout the genome and di-methylation is loosely linked with repressed genes, though this is poorly defined in most cell types. Tri-methylation of H3K27 is closely associated with repressed genes, presumably due to binding of the CBX family of proteins contained in the PRC1 complex to K27me3 and subsequent repression by this family of complexes (Bernstein et al., 2006). We performed western blots on whole cell lysates from WT, knockout and the point mutant cells and probed them with antibodies to di- and tri-H3K27 methylation and EZH2 (Fig 2C and Supp. Fig. 3). H3K27me3 was reduced to levels we could not detect in the mutant cells, as was seen in the knockout cells, but the levels of di-methylation were largely unaffected in the R681C mutant cells.

The mutant ES cells look phenotypically similar to WT cells and self-renew (Fig 2D). This was the anticipated result since knockout cells in the PRC2 core components EZH2 and EED are also phenotypically similar to WT cells and can self-renew. We examined whether the mutant cell lines would form embryoid bodies similar to those formed by WT cells, would apoptose like *Ezh2* or *Eed* knockout cells, or would take an alternate path. We formed embryoid bodies by the hanging drop procedure and compared the resultant phenotypes of WT, the point mutant and knockout cells. We found that the mutant cells formed embryoid bodies, making them phenotypically separate from the knockout cells (Fig 2D). This revealed that the dramatic reduction in H3K27me3 we see in the point mutant cells does not stop the cells from differentiating, at least at a gross level.

We determined whether the mutant PRC2 complex was localized to the same genomic regions as the WT complex by characterizing peaks of the core PRC2 component SUZ12 through use of CUT&RUN. There were fewer peaks of PRC2 chromatin binding in the mutant cells when compared to WT, as expected due to the lack of H3K27me3 which facilitates binding and spreading of PRC2 (Oksuz et al., 2018). There was a high degree of overlap between peaks of occupancy in ES cells (Fig. 2E). We also observed overlap in binding of SUZ12 between WT and mutant cells at Day 4 and Day 8 of differentiation into embryoid bodies (Fig. 2F). We conclude that in these cells the major effect of the point mutation was to reduce the tri-methylation of H3K27. This is similar to the results found in other studies of *Ezh2* point mutations (Lavarone et al., 2019).

To further test the developmental potential of the 681C mutant embryonic stem cells we assessed the formation of beating clusters of cardiac cells. For this assay cells are differentiated in hanging drops and then plated in individual wells to form a monolayer in differentiation media. After 10-15 days of differentiation each well is visually examined for the presence of pulsing cells. Beating clusters formed from both WT and mutant cells. There was some variation in the proportion of EBs that could form beating clusters between the two 681C mutant strains, but in all cases the mutant cells could form beating clusters (Fig. 2G). These data bolster the conclusion that the reduction in H3K27me3 does not prevent cells from differentiating. The R681C mutation therefore offers an opportunity to examine the contribution of tri-methylated H3K27 to gene regulation during differentiation and raised the issue of whether molecular changes are caused by the point mutation.

### Molecular phenotype of *Ezh2* point mutant cells

To determine whether there were molecular phenotypes associated with the hypomorphic mutation of *Ezh2*, and the resultant loss of H3K27me3, we determined the target genes of PRC2 in our cell lines and measured their gene expression levels. To annotate target genes in this system, we performed CUT&RUN analysis using antibodies to H3K27me3, RING1b, EZH2 and SUZ12 in WT cells as embryonic stem cells (ESC) and as Embryoid bodies after differentiation for eight days (D8EB). Target genes were defined as having a broad peak of H3K27me3 over the transcription start site in at least two time points. There have only been a small number of studies of PRC2 localization in differentiating cells, but of the approximately seven thousand target genes we identified, just over 70% overlapped with previously published data (Fig. 3A and (Juan et al., 2016)). We note that peaks called from our data set that did not overlap with previous called peaks showed signal in the published data that was below the threshold, indicating further agreement between the analysis done here and previous work. There were far more targets called using the H3K27me3 than with the individual components of either polycomb complex, though the targets are largely overlapping (Fig. 3B). This is likely due to differences in the strength of individual antibodies. Therefore, we used the targets identified with H3K27me3 for further analysis. In keeping with the western blot from whole cell lysates, H3K27me3 was reduced on target genes in mutant cells (Fig. 3C). Levels of H3K27me3 were significantly lower in mutant cells at day 4 of differentiation and then were increased at day 8 of differentiation but to levels well below WT levels. The residual tri-methylation at day 8 might be due to EZH1 function (Lavarone et al., 2019; Margueron et al., 2008; Shen et al., 2008). We focus below on the events that happen during these first four days of differentiation and the impact of the lack of H3K27me3 during this time frame. We conclude that the ability of the mutant cells to differentiate was not due to retention of normal tri-methylation levels specifically on target genes.

**Figure 3.**
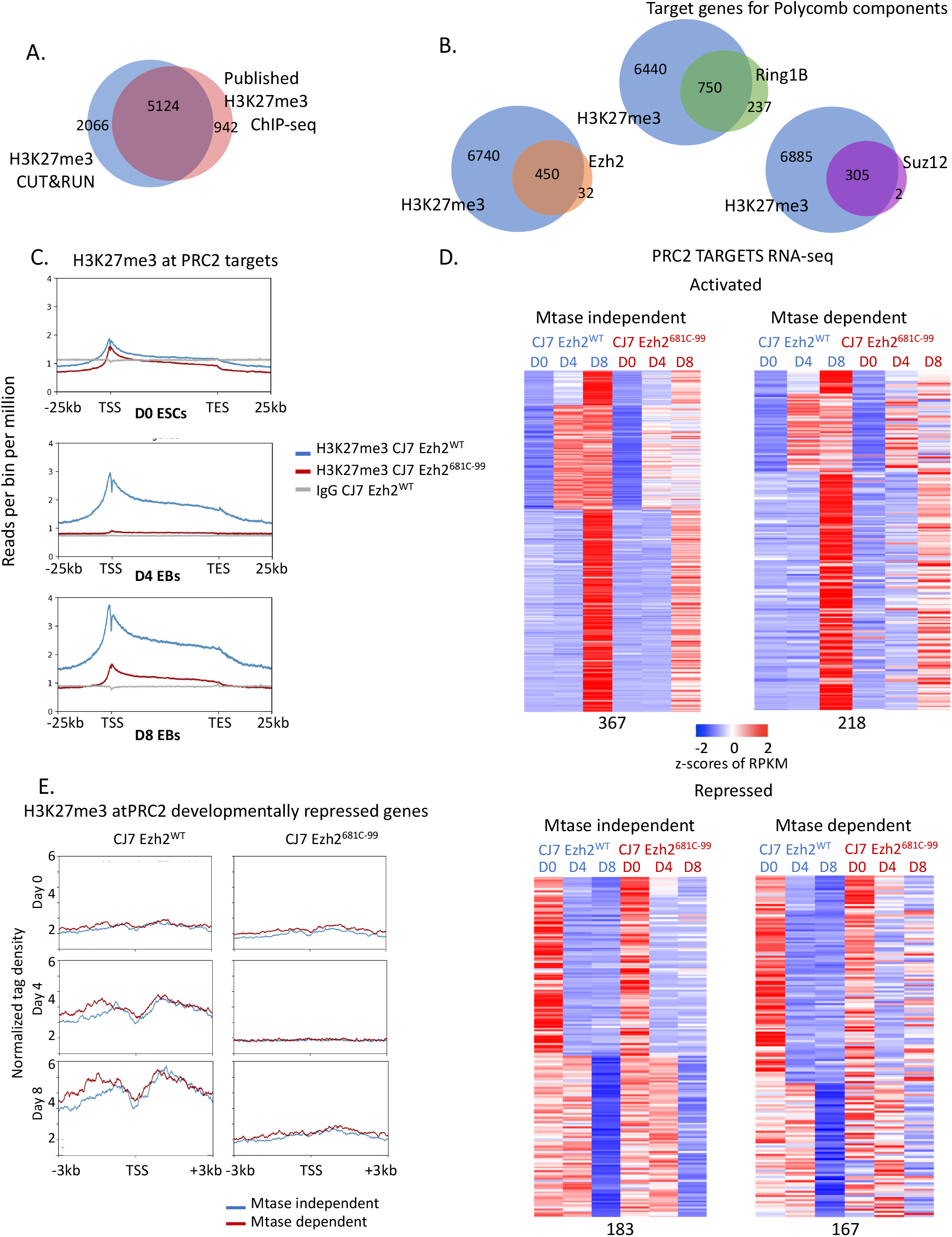
(A) Venn diagram showing overlap of H3K27me3 target genes assigned from CUT&RUN peaks from WT cells and published ChIP-seq data peaks. (B) Venn diagrams showing overlap of target genes of H3K27me3 and other PRC2 components by CUT&RUN in WT cells. (C) Metaplots over PRC2 target genes showing H3K27me3 CUT&RUN versus IgG control signal at Day 0, 4 and 8 of embryoid body formation in wild-type and mutant cells. (D) Heatmap showing expression patterns of Mtase dependent and independent genes in wild-type and mutant ES cells over developmental time-course. Z-scores of average RPKMs of triplicates are shown for each gene across all samples. Genes are separated based on if they were activated (left two panels) or repressed (right two panels) in wild-type cells over time as well as if they had a similar expression pattern in mutant cells (Mtase dependent genes) or a different expression pattern (Mtase independent genes). The four groups are further clustered based on fold-changes over time in untreated cells. (E) Metaplots showing H3K27me3 binding in wild-type and mutant cells by cut- and-run at Mtase dependent or independent and developmentally repressed PRC2 target genes over the time course.

The experiments described above using small molecule inhibitors revealed that there are genes that require H3K27me3 for their regulation and those that do not require H3K27me3 to maintain proper regulation. We investigated whether the point mutant cells showed these same two gene classes. We performed RNA-seq analysis from ES cells and both D4 and D8EBs from WT, CJ7 Ezh2^681C-99^ and CJ7 Ezh2^681C-102^ cells. The data for both mutant cell lines are nearly overlapping and so for clarity we present the data from the CJ7 Ezh2^681C-99^ (Supp. Fig. 4). For this analysis we defined PRC2 target genes as those that had a statistically significant peak of H3K27me3 from CUT&RUN data at either D4 or D8 during the EB formation protocol. PRC2 target genes were identified using wild-type H3K27me3 peaks called by Homer using a p-value threshold of 0.001, a length of at least 1500 bp, 1rpkm in at least two time points and signal overlapping Refseq annotated genes (TSS+-5kb). As with the cells treated with the small molecule inhibitor, we could separate genes that rely on H3K27me3 from those that do not need a high level of the modification for their regulation (Fig 3D). The gene expression patterns from the mutant cells and from the GSK343 treated cells clustered based upon day of differentiation rather than by whether they were drug treated or mutant (Supp. Fig. 5), although the correlation was not as strong between drug treated cells and mutant cells as that seen between the two mutant cell lines. We detected both activated and repressed genes that fell into methyltransferase-dependent and –independent categories. The 350 genes that are normally repressed during differentiation in WT cells, and were either similarly repressed or were not repressed in mutant cells, showed more consistent patterns than the genes that were normally activated in WT cells. These patterns were well established by D4 of differentiation, thus we used repressed genes to further analyze any characteristics specific to the methyltransferase dependent or to the independent genes.

We examined whether the level of H3K27me3 normally found on the genes in WT cells or the amount of signal remaining in the mutant cells could predict whether a gene would be dependent on the modification for its regulation. However, there was not a difference in the levels of tri-methylation based on whether the genes require this modification for their regulation (Fig. 3E). There was also strong overlap in the genes with PRC2 peaks suggesting that the complex was still targeted normally (Supp. Fig 6) We then examined several characteristics of 350 genes that are normally repressed during differentiation of WT ES cells including other histone modifications, CpG methylation, and further sub dividing genes by their dependence on H3K27me3 (Supp. Fig. 6). None of these characteristics showed any significant differences between genes who repression was dependent upon methyltransferase activity and genes whose repression was not dependent on methyltransferase activity. We conclude that there is a variable reliance on H3K27me3 for gene regulation and that H3K27me3 is not the only driver of PRC2 target gene repression during EB formation, just as full levels are not needed for differentiation.

### Methyltransferase point mutants cannot maintain differentiated cell identity

A major issue in developmental gene expression concerns the interplay between establishment and maintenance of gene expression profiles during differentiation. The Polycomb group (PcG) system, including PRC2, plays a role in both aspects of gene regulation in flies and in mammals. Substantial early work on mutant Drosophila highlighted a role for the PcG, including gene products now known to compose PRC2, in maintenance. Given that we saw limited impact of the ablation of H3K27me3 on establishment of the differentiated EB phenotype, we tested whether the mutant or drug treated mouse cells were able to maintain a differentiated state.

Under normal conditions, cells differentiated into embryoid bodies cannot revert to ESCs without major manipulation such as introducing Yamanaka transcription factors (Nakagawa et al., 2008; Takahashi & Yamanaka, 2006). To determine if the 681C point mutant cells stably committed to a differentiated state, we formed day 8 EBs with WT and mutant cells, dissociated the embryoid bodies into single cells and re-plated them in ESC conditions without any additional manipulation. The cells were cultured in the ESC media for five days and then stained for alkaline phosphatase (AP) activity. After incubation with an appropriate colorometric substrate, ESCs become bright pink due to their alkaline phosphatase activity, while feeder cells or any other differentiated cells remain unstained. We found a ten-fold increase in the number of AP-positive colonies generated by the mutant cells over the WT (Fig. 4A and 4B). We asked whether this lack of commitment was maintained after longer periods of differentiation and found that after 14 days there remained more cells that could revert to ES cells, but to a considerably lesser extent than seen after 8 days (Figs. 4A, 4B). We conclude that the mutant cells are not stably committed to the more defined lineages but can switch back to an undifferentiated state, but that this flexibility decreases after two weeks of differentiation.

**Figure 4.**
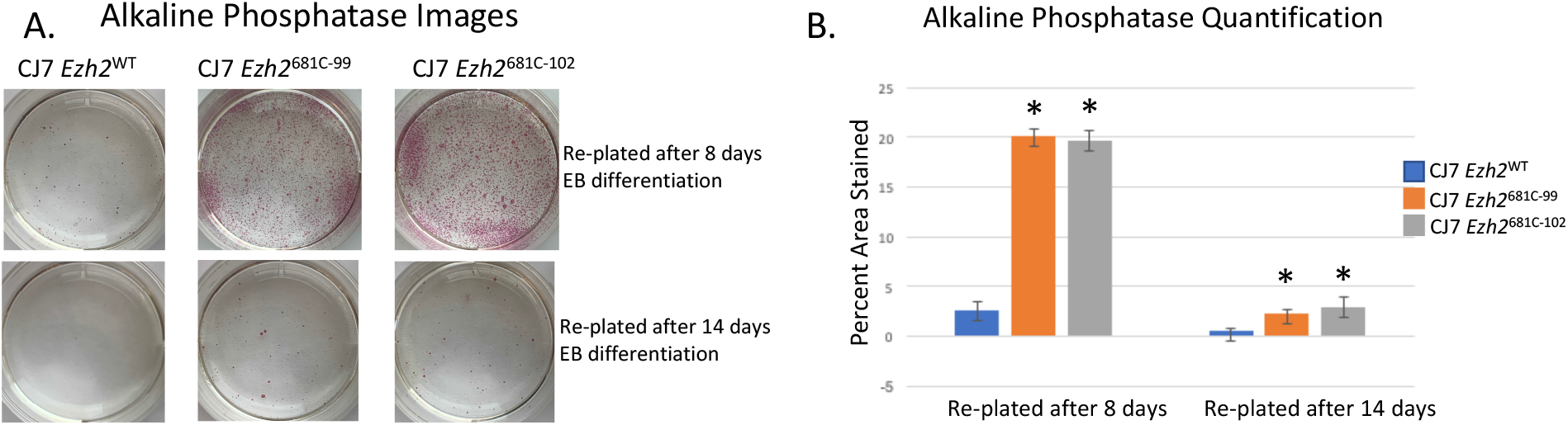
(A) Alkaline phosphatase staining from WT, 681C-99 and 681C-102 cells that had been differentiated for 8 or 14 days and then transferred into ESC conditions for five days before staining. ESCs stain bright pink in this assay. (B) Quantification of alkaline phosphatase staining from re-plated cells with standard deviation error bars. Samples with significant difference (p<0.05) from WT cells marked with an asterisk.

We used the small molecule inhibitor GSK343 to examine the time period during which this flexibility is established. There were three possibilities: a) full levels of H3K27me3 might be needed continuously to maintain the differentiated state; b) they might be needed at specific times as the cells are differentiating; c) they might be needed only when cells are challenged with external stimuli that allow reversion to embryonic stem cells. Examination of the mutant cells addressed blocking full levels of methylation throughout the experimental time course, but did not address the time period when that lack of activity was most important. To separate the possibilities, we added or removed GSK343 during the differentiation time-course (Fig. 5A shows the experimental design). To mimic the WT and point mutant conditions a subset of cells were treated continuously with DMSO or GSK343 respectively. These treatments served as both a baseline for altered times of GSK343 application. They also served to validate that the results observed with re-plating of mutant cells were not caused by off target effects of the CRISPR genetic manipulation or by mutations acquired during selection of these cells. Cells continuously treated with GSK343 had higher numbers of AP-positive cells following re-plating at 8 days, similar to the mutant cells; in contrast to the mutant cells there even more AP-positive cells following re-plating after 14 days of differentiation in GSK343. GSK343 treatment affects both di- and tri-methylation which might account for the increased plasticity seen at later time points in drug treated cells (Supp. Fig. 7).

**Figure 5.**
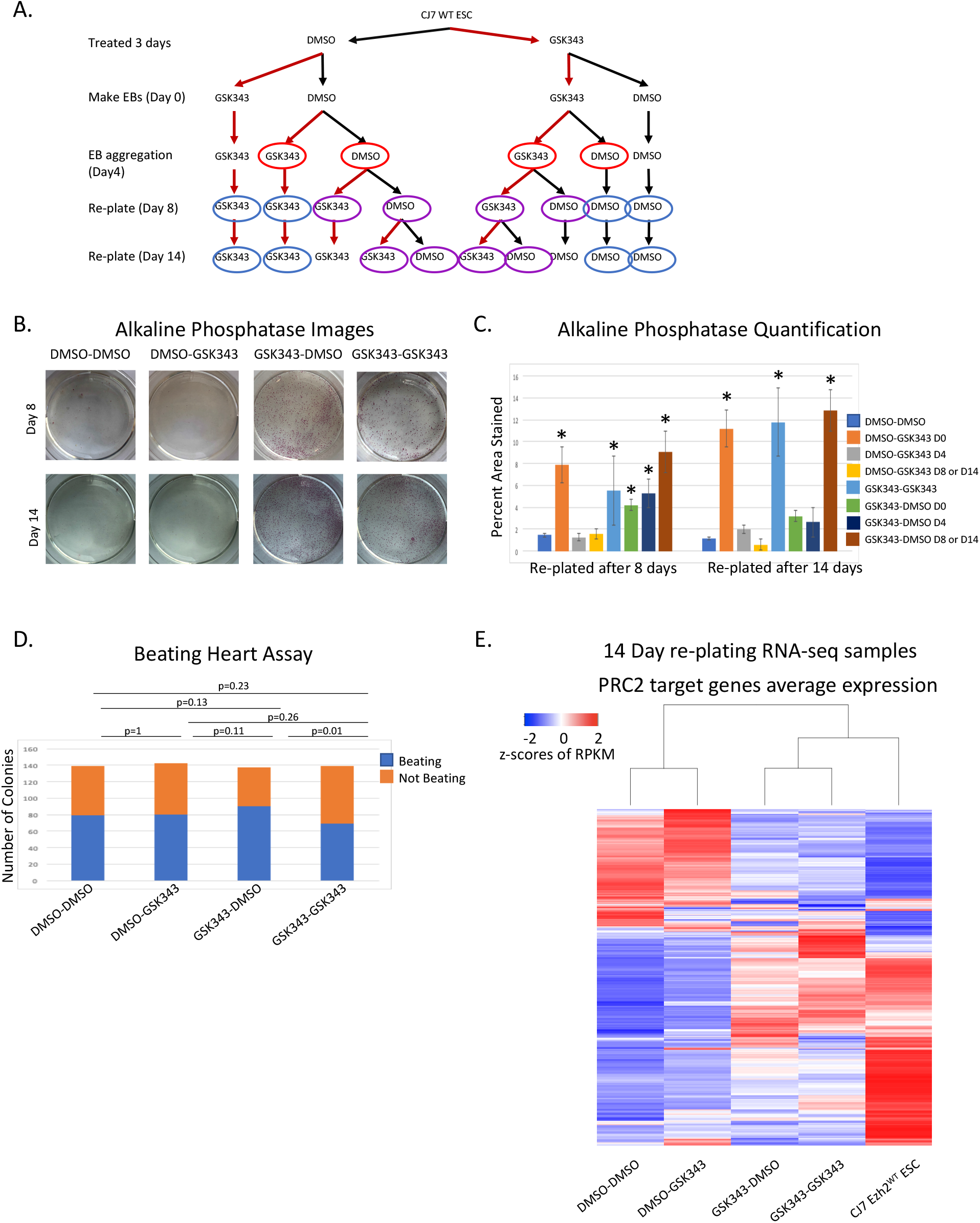
(A) Diagram showing the drug treatment plan for differentiation and re-plating of cells. Red circles indicate the cells used for the beating heart assay in panel D, Purple are shown in images in (B) as well as quantified in (C) and blue circles highlight samples only quantified in (C). (B) Alkaline phosphatase staining from treated cells re-plated after 8 or 14 days of EB differentiation. Only cells that start in GSK343 treatment stain pink indicating that the cells have reverted to an ES phenotype. (C) Quantification of alkaline phosphatase staining from cells treated with DMSO or GSK343. Time of the treatment switch is indicated in the legend. Samples with significant differences from those continuously treated with DMSO (p<0.05) are marked with an asterisk. (D) Quantification of beating heart assay with GSK343 and DMSO treated cells. Cells were differentiated as EBs for 4 days and then plated into differentiation media. Blue indicates the number of EBs that formed beating cells and orange indicates the number that did not. (E) Unsupervised clustering of RNA-seq average expression of cells re-plated for 5 days after 14 days of EB formation. Cells that were treated with GSK343 have gene expression profiles that cluster with average expression from WT ESCs.

In addition to continual treatment with inhibitor, we varied the timing of GSK343 addition as depicted in Fig. 5A. Briefly, we treated ES cells for three days with either DMSO or GSK343 before starting the differentiation. When embryoid body formation was initiated or at day 4 of differentiation we changed the treatment from DMSO to GSK343 (or vice versa) for half of the cells. We also switched the treatment at the time of re-plating into the ES conditions at day 8. Finally, we allowed cells to differentiate to day 14 and switched treatment regimen (Fig. 5A). The treatment type was switched just once in each scheme so we could determine whether blocking the methylation of H3K27 at early or later stages of differentiation had the greatest effect. These experiments allow us to determine whether reducing H3K27me3 has the greatest effect on cell identity when the cells are differentiating, when they are challenged by re-plating or throughout the differentiating time-course.

We found that the crucial window for treatment with the inhibitor was in the first four days of embryoid body formation (Fig. 5A, B, C). Increased staining by alkaline phosphatase activity was seen in all cells that had initially been treated with inhibitor, regardless of when it was removed (Fig. 5C). Notably, treating with GSK343 for three days prior to inducing differentiation, then removing the inhibitor when differentiation was initiated, led to a significant increase in ES cells following re-plating at day 8. The only case where we saw increased staining in re-plated cells that were initially treated with DMSO was when the inhibitor was added at the onset of differentiation (Fig 5B, C). We conclude that the inhibition of methyltransferase activity either immediately prior to differentiation or during the first four days of differentiation allows the cells to remain in a more plastic state such that they can revert to a stem cell-like phenotype when placed in the proper growth conditions.

When we examined the ability of treated cells to form beating colonies after treatment with the small molecule we found no difference in the proportion of colonies that spontaneously start beating between any of the treatment groups (Fig. 5D). This is consistent with the data from the point mutant cells where both WT and mutant cells could form beating colonies and further indicates that there is no deficit in the ability of the cells to differentiate when H3K27 methylation is inhibited. Thus, as was seen with the comparison of WT and point mutant cells, the obvious difference between DMSO and small molecule treated cells occurred when cells were challenged to grow in the ESC culture conditions (Figs 5B,C). We conclude that H3K27me3 methylation is more important in maintaining the differentiated state than in generating that state, and that the critical time window occurs early in the differentiation process.

The GSK343 treated cells and the mutant cells both appeared to revert to a pluripotent state. To verify that these cells retained developmental potential we re-differentiated these cells and determined whether they display the characteristic ability of ES cells to develop more committed cells. We used the re-plated cells from both 681C mutant cells and those treated with the inhibitor and attempted to make embryoid bodies. Both sets of re-plated cells were able to make embryoid bodies, which confirms that they have developmental potential (Data not shown).

To examine whether the reversion to a pluripotent state involved substantive changes in gene expression, as opposed to gene expression changes in a few key genes, we examined genome-wide gene expression pattern of the reverted cells and compared those to ESCs. To examine the molecular phenotype of the re-plated cells, we isolated and sequenced RNA from drug treated day 14 cells that were re-plated and had reverted to ES cell phenotype. Unsupervised clustering showed that the average expression profiles of the cells that had been treated with GSK343, and therefore had much higher levels of reversion, were more like WT ES cells than to the DMSO treated and re-plated control cells (Fig. 5E.) We conclude that the cells we see staining with alkaline phosphatase in our re-plating assays are reverting to an ES phenotype. We examined the subset of PRC2 target genes that are normally repressed during differentiation and found that many are reactivated when cells treated with GSK343 are re-plated. This differs from the genes that are reactivated in the cells initially treated with DMSO. The changes in gene expression observed following re-plating of cells treated with GSK343 were significantly different from the patterns observed after re-plating of cells treated with DMSO. (Supp. Fig. 7C). These expression pattern changes show the same response to the time of treatment with GSK343 as seen above; reversion to the WT pattern requires GSK343 treatment early in differentiation. From these data, we conclude that though all cells are placed under developmental stress when re-plated into ES conditions, only those where H3K27 methylation has been blocked revert to an ES cell phenotype. This underlines the importance of K3K27me3 in establishing the heritable gene expression profile of differentiated lineages.

## Discussion

These studies offer two advances in understanding the role for tri-methylation of H3K27 during differentiation of ES cells into embryoid bodies. First, many PRC2 targets continue to be regulated in a normal manner during the first four days of differentiation despite significantly reduced H3K27me3 levels on these targets (Fig. 3.) Thus, H3K27me3 is not necessary for repression of a significant set of PRC2 targets, indicating compensating mechanisms for repression of these genes. Second, while many PRC2 targets are dysregulated when H3K27me3 is blocked, we did not observe a significant impact on differentiation. In contrast, there was a large enhancement of the ability of differentiated cells to revert to a pluripotent phenotype when placed into embryonic stem cell culture conditions. We conclude that the primary role for H3K27me3 during early differentiation is maintenance of the differentiated state.

Differentiated WT cells are not generally capable of reverting to an ES phenotype when their growth conditions are altered by a change in media. In normal cases, it takes the reactivation of key transcription factors to allow cells to return to that state. In contrast, *Ezh2* point mutant cells and those that that have been treated with a small molecule inhibitor readily revert to an ES phenotype when placed into media that supports that type of growth. This was seen at both the cellular and molecular level. When we varied the time windows where inhibitor was present, we found that the crucial window for normal H3K27me3 levels was between the start of differentiation through the first four days of embryoid body formation. Not having the ability to add the tri-methylation modification at those early stages of differentiation sets the stage for the cells to be able to revert to an ES phenotype when challenged, even if H3K27me3 is restored later during embryoid body formation. In the setting of the early stages of ES differentiation into embryoid bodies, the H3K27me3 modification is acting analogously to a stopper on a swinging door. When the modification is present the door will only open one way and the cells cannot go backward, but removal of the modification enables the door to swing both ways allowing cells to go back and forth between the differentiated and pluripotent state in response to external signals.

There are potential implications for these data in terms of the use of small molecule inhibitors in therapeutic situations. Multiple adult cell types have different requirements for PRC2 during their differentiation. Most relevant to the inhibition of PRC2 is the development of blood cells. PRC2 is required for the differentiation of blood stem cells and is the site of some of the highest levels of PRC2 expression in healthy adult tissues. If the phenotypes that we have observed during embryoid body formation occur in a similar manner in blood stem cells, treating patients with the small molecule inhibitors might alter the stability of commitment of healthy stem cells, raising the possibility of novel cancers arising from cells that cannot stably differentiate. Indeed, at least one clinical trial was temporarily suspended because patients had developed novel cancers (Fioravanti et al., 2018; Harris, 2018; Italiano et al., 2018). Determining the effect of blocking H3K27me3 in other differentiating cell lineages might be important to clinical intervention by expanding the knowledge of the potential side effects.

We note that it is difficult to completely eliminate H3K27 methylation with mutations (Lavarone et al., 2019), and that the mutation we generated in *Ezh2* specifically impacts H3K27me3, especially early in differentiation, but does not eliminate all methylation of H3K27. This is both a limitation and an advantage; the significant impact on H3K27me3 early in differentiation allowed us to show that loss of tri-methylation has a potent phenotype (unstable commitment) at this stage yet does not have a discernible differentiation phenotype. These data demonstrate a striking difference in the dependency of these two key phenotypes on H3K27me3. This *Ezh2* mutation also has a molecular phenotype when gene regulation is examined, raising the possibility that the network of genes that require H3K27me3 for appropriate regulation at this stage are primarily responsible for driving stable commitment to the differentiated state.

## Methods

### Cell Culture

ES cells: CJ7 WT, CJ7 *Ezh2-/-* and CJ7 *Eed-/-* cells were a generous gift from the laboratory of Stuart Orkin. We cultured all embryonic stem cells on a monolayer of feeder MEFs in ES media (DMEM (Gibco 11995-506) supplemented with 20%FBS, NEAA, pen/strep, glutamax, and LIF) on tissue culture treated flasks coated with 0.2% geletin. Media was changed daily and cells were split every 2-3 days.

EZH2 point mutant Cells: Point mutations were introduced into CJ7 WT using CRISPR RNP transfected into cells with the Amaxa mouse ES kit (Lonza VPH-1001). 1×10^6^ cells were transfected with the RNP containing two separate guide RNAs a single stranded donor oligo and a linearized puromycin resistance gene. Cells were plated into three wells of a six well plate with puroR MEFs and regular ES media. The next day puromycin was added to the wells in a range of concentrations and kept on the cells for the next two days. Following selection, resistant cells were expanded, individual colonies were then re-plated into twenty-four well plates and checked for the presence of the mutation by restriction enzyme digestion followed by confirmation with sequencing.

EB formation: Embryoid bodies were formed using the hanging drop method(Behringer et al., 2016; Dang et al., 2002). Briefly, ES cells were trypsinized, de-MEFed and then resuspended in differentiation media (IMDM (Gibco 12440-053) supplemented with 20% FBS, pen/strep, and glutamax. No LIF). Droplets containing approximately 180 cells were incubated for four days so that spheres of differentiating cells could form. After the four days EBs were collected and grown in suspension in non-adherent plates or used for subsequent experiments.

Beating heart assay: Beating clusters were differentiated from day 4 embryoid bodies (Boheler et al., 2002; Hescheler et al., 1997). Individual EBs were transferred into individual wells of a gelatinized 12 well plate. Cells were maintained in differentiation media throughout the experiment. Wells were monitored for beating colonies for the next 15 days and were scored as positive if there were any beating cells in that time period.

### Western Blot

The antibodies that were used probe the Western blots were from Cell Signaling Technology H3K27me3 (9733S) and H3K27me2 (9728S) as well as from EMD Millipore EZH2 (07-689).

### RNA-seq

RNA was isolated from whole cells using the Nucleospin RNA kit from Macherey Nagel (740955.50). We then depleted rRNA using the Ribozero Gold kit from Epicentre (RZG1224). cDNA was synthesized from the purified RNA using the Superscript Vilo cDNA sequencing kit from ThermoFisher (11754050) the libraries were assembled as previously described (Bowman et al., 2013). Three replicates were completed for all experimental conditions except the Day 14 re-plated cells where two replicates were completed.

### Methyltransferase Assay

Purified protein complexes were incubated at room temperature with a histone substrate at a final concentration of 500nM and approximately 500nM radioactive SAM (Adenosyl-L-methionine S-methyl^3^H from PerkinElmer (NET155H250UC)) in methyltransferase buffer (10% glycerol, 25mM HEPES PH 7.9, 2mM MgCl with 1mM DTT added fresh). Unless otherwise specified all reactions ran for one hour before being stopped with the addition of 6x SDS buffer. Samples were then run on an SDS page gel, coomassie stained and incubated in AMPLIFY Amersham/GE (NAMP100) for 20 minutes. The gel was dried and exposed to film for a minimum of 24 hours before developing.

### Protein purification

The ORF for each member of the core PRC2 complex were cloned into the pFastBac1 baculovirus with Suz12 tagged with 3xFlag. Sf9 transfection with bacmid DNA and virus amplification were performed essentially as described for the Bac-to-Bac Baculovirus Expression System (ThermoFisher Scientific). Sf9 cells were maintained in Hyclone CCM3 (CCM3 liquid medium with L-glutamine, GE Healthcare Life Sciences SH30065.02) supplemented with 50 U/ml Penicillin-Streptomycin (ThermoFisher Scientific, 15140-122). For protein expression, 2×10^6^ Sf9 cells/ml were infected at an MOI of approximately 10. Cells were harvested 66 hours post infection by centrifugation at 5000xG for 15 minutes. Cells were lysed, treated with DNAse1 then the complex was bound to M2 and the complex was eluted with flag peptide.

### CUT&RUN

CUT&RUN experiments were performed as described (Skene & Henikoff, 2017). The antibodies we used were from Cell Signaling Technology H3K27me3 (9733S), EZH2 (5246S), SUZ12 (3737S) and Bethyl laboratories RING1b (A302-869A). Libraries were constructed as described in the RNA-seq section. Two replicates were completed and representative experiments are shown.

### Alkaline Phosphotase Assay

Cells were stained according to the instructions in the Stemgent AP Staining Kit II from Reprocell (00-0055). Images of stained wells were quantified using ImageJ FIJI software.

### Bioinformatics Methods

RNA-seq data processing: All RNA sequencing reads were aligned to the mm10 genome using STAR v2.5.3 (Dobin et al., 2013). Gene annotations were obtained from Ensembl (Hunt et al., 2018) Genome browser tracks were generated using Homer v4.10.3 (Heinz et al., 2010) and visualized in IGV (Robinson et al., 2011). Reads in exons of Refseq annotated genes were counted using featureCounts v1.6.1(Liao et al., 2014). edgeR (Robinson et al., 2010) was used to normalize reads and calculate FDR using triplicates. All further calculations and figures were made using R v3.3.2 (Dessau & Pipper, 2008). For all heatmaps, standard z-scores calculations were made using average RPKMs of triplicates for each gene across all conditions. Genes were narrowed down to only PRC2 targets using H3K27me3 CUT&RUN data. Only genes changing in untreated or wild-type cells over the time-course were used for further analysis. The genes were filtered using cut-offs of at least 0.5 RPKM average expression at any timepoint, at least 1.5 fold-change and maximum 0.05 FDR while comparing Day 8 and Day 4 or Day 4 and Day 0. Genes were then clustered based on up or down-regulation in control cells and up or down-regulation or unchanged gene expression in mutant or treated cells.

For Fig. 5E, unsupervised clustering was performed using R’s heatmap.2 function. CpG and GC levels were calculated using Homer’s annotatePeaks function.

CUT&RUN data processing: All CUT&RUN reads were aligned to the mm10 genome using bowtie2 and filtered using samtools (Li et al., 2009) to keep uniquely aligned reads. Genome browser tracks were generated using Homer v4.10.3 and visualized in IGV. Peaks were called using Homer’s findPeaks function for broad peaks. PRC2 target genes were identified using wild-type H3K27me3 peaks at least 1500 bp in length, 1rpkm in at least two time points and overlapping Refseq annotated genes (TSS+-5kb). Target genes for Suz12 and Ezh2 were identified using the same parameters as H3K27me3. Average of RPKMs of two replicates was calculated for Suz12 CUT&RUN data, and target genes were identified using the same parameters as H3K27me3. For Figures 2E and 2F, average of RPKMs of two replicates was used to find Suz12 peaks. For Figure 2F, peaks from D4 and D8 were merged. Deeptools v3.3.0(Ramírez et al., 2014) was used to make average profile plots for Fig. 3E.

Published ChIP-seq data sources and processing: Published data for other canonical and non-canonical PRC2 components was downloaded from GEO as follows : H3K27me3 (GSM2282188, GSM2282191, GSM2282192) (Juan et al., 2016), EPOP (GSM2098943) (Beringer et al., 2016), Jarid2 (GSM491760) (Li et al., 2010), H3K4me1 (GSM1180178), H3K4me3 (GSM1180179), H3K9me3 (GSM1180180), H3K36me3 (GSM1180183) (Hon et al., 2014). FASTQ files were downloaded from GEO using sratools v2.9.1 (https://ncbi.github.io/sra-tools/). All reads were aligned to the mm10 genome using bowtie2 and filtered using samtools to keep uniquely aligned reads. Genome browser tracks were generated using Homer v4.10.3. Deeptools v3.3.0 was used to make average profile plots for Fig. 3C and E.

## Supplemental Figure Legends

**Supp. Fig. 1.**
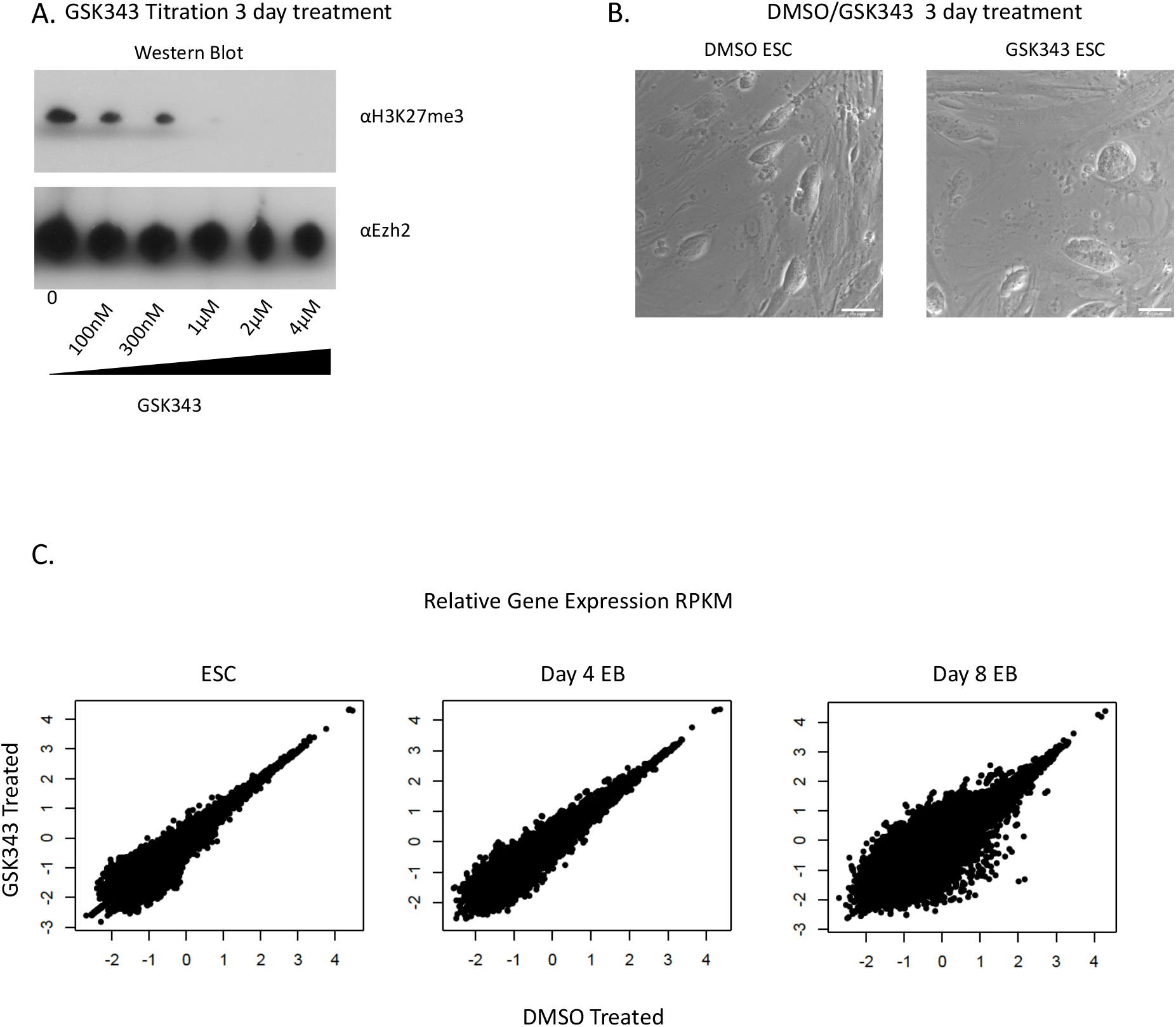
(A) Western blot of WT CJ7 cells treated for three days with an increasing concentration of GSK343 probed with antibodies to H3K27me3 and EZH2. The 4uM concentration was non-toxic to cells and yielded a loss of H3K27me3 signal and so this concentration was used for all experiments. (B) 54x magnification of ES cells treated for three days with DMSO or GSK343. ES colonies show characteristic halo and are indistinguishable between treatments. (C) Comparisons of normalized gene expression (RPKM) from cells treated with DMSO or GSK343 and differentiated as embryoid bodies. At the genome level there are many genes that have the same expression pattern across the two treatments.

**Supp. Fig. 2.**
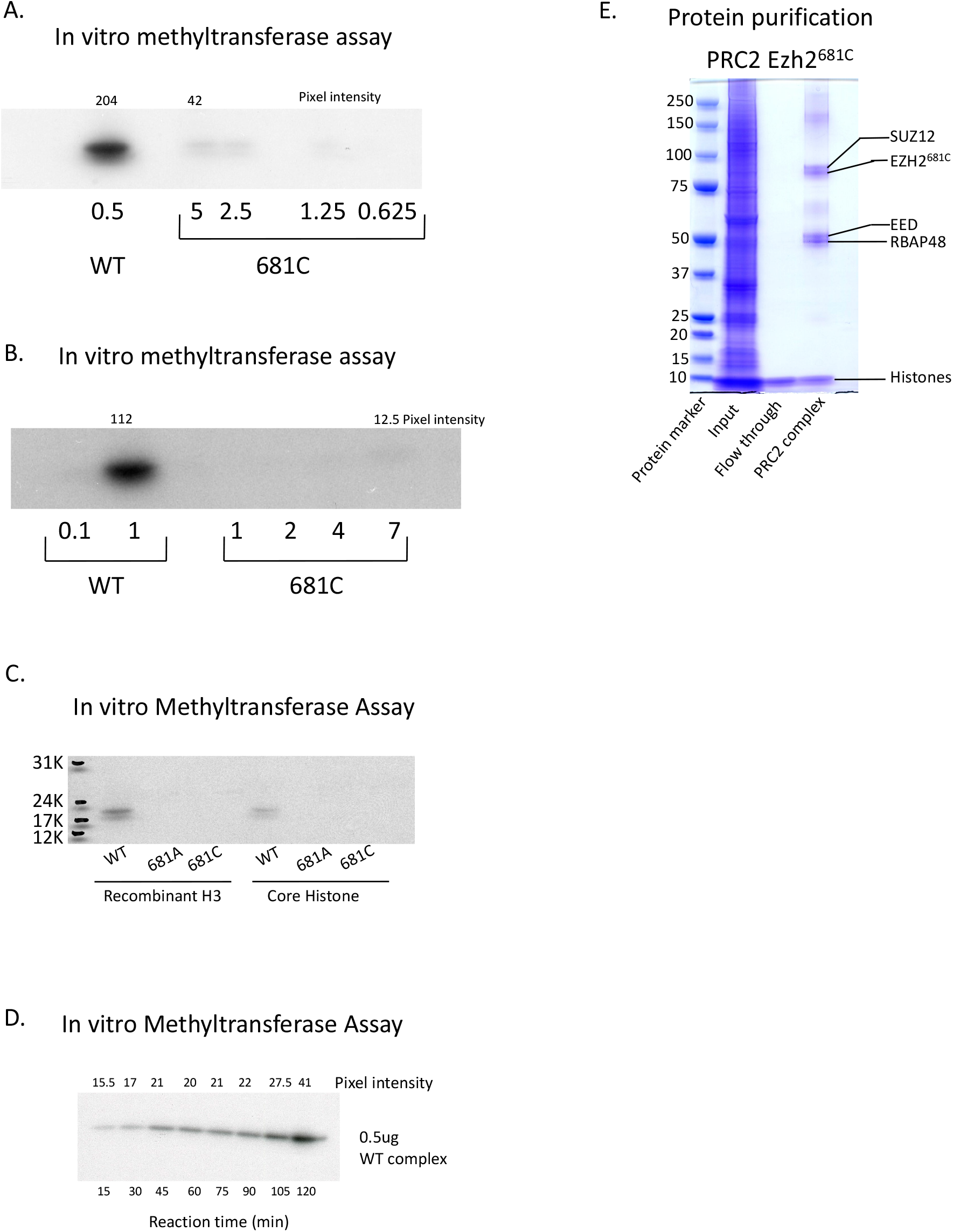
(A,B) In vitro methyltransferase assay comparing EZH2-WT and EZH2-681C. Concentration of the PRC2 complex is indicated below the image with pixel intensity indicated above. (C) In vitro methyltransferase assay with WT, 681A and 681C complexes with both recombinant histones that do not have any covalent modifications and core histones that do as the substrate. With both substrates, the mutant complexes did not show signal. (D) In vitro methyltransferase assay with WT complex over time. All other methyltransferase assays shown here were done for 60 minutes which is well within the linear range. (E) Coomassie stain of mutant PRC2 complex. Complex was purified from Sf9 cells by using a Flag-tagged SUZ12 and purifying over flag beads.

**Supp. Fig. 3.**
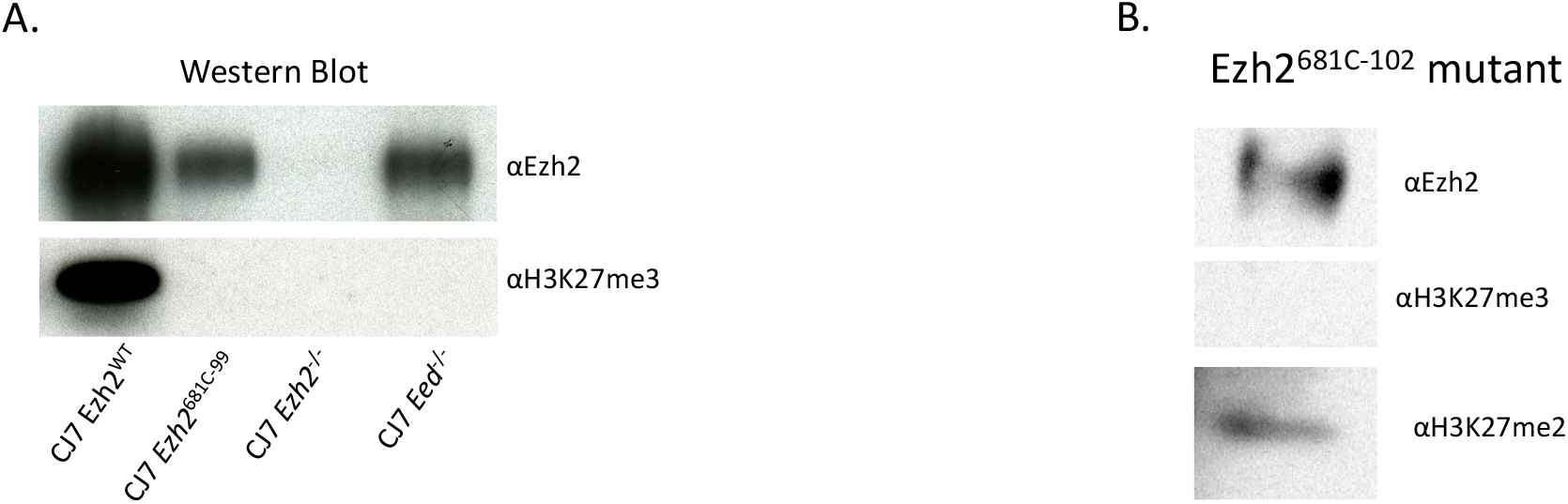
(A) Western blot showing the EZH2 and H3K27me3 signal for WT, EZH2 mutant and PRC2 KO cells. (B) Western blot showing the EZH2, H3K27me3 and H3K27me2 signal of EZH2^618C-102^ mutant cells.

**Supp. Fig. 4.**
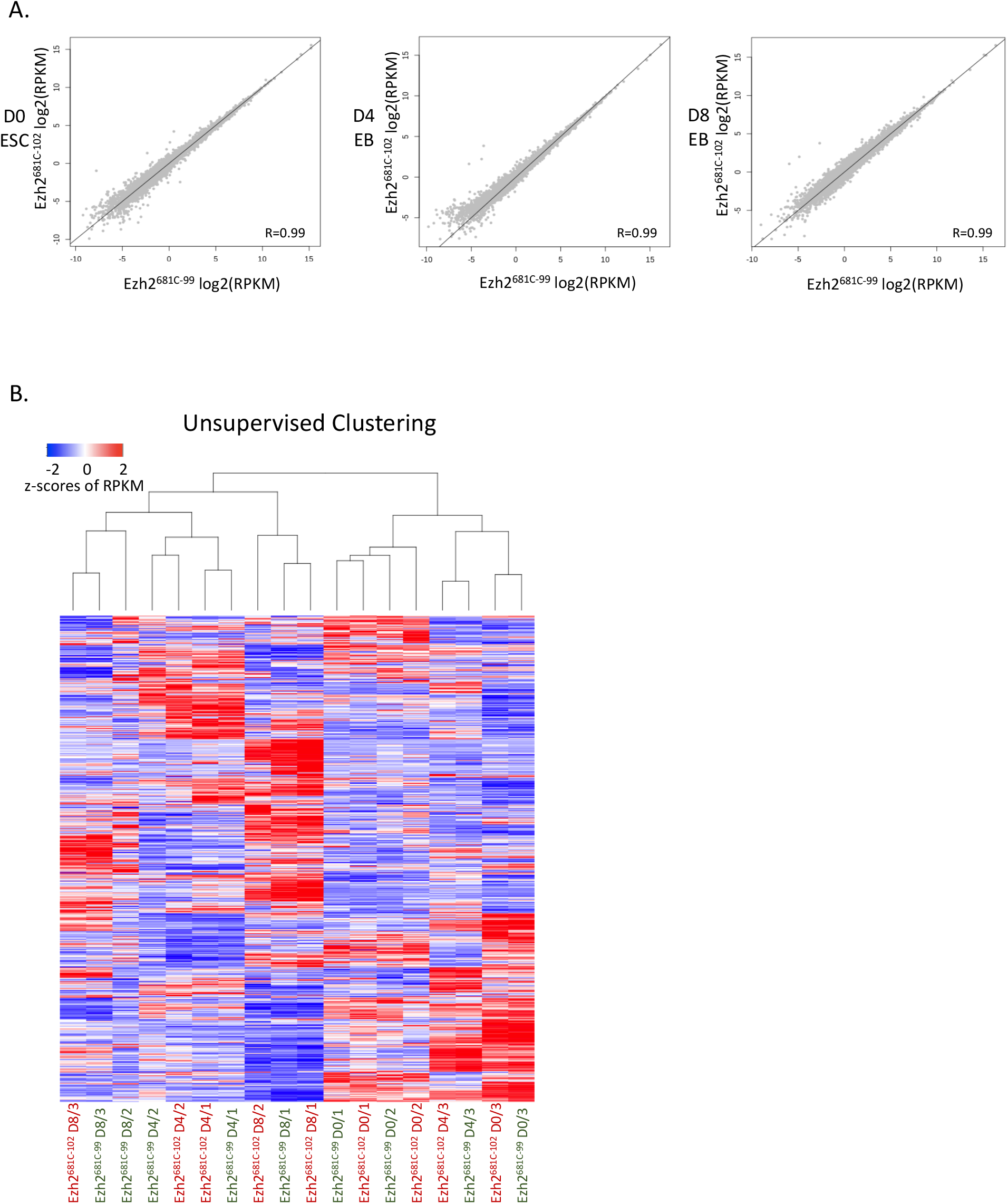
(A) Comparisons between Ezh2^618C-99^ and Ezh2^681C-102^ RNA-seq over the differentiation time-course. The two mutants are nearly identical at all time points. (B) Unsupervised clustering of the mutant RNA-seq showing that the day of differentiation is more relevant predictor of similarity rather than the mutant strain.

**Supp. Fig. 5.**
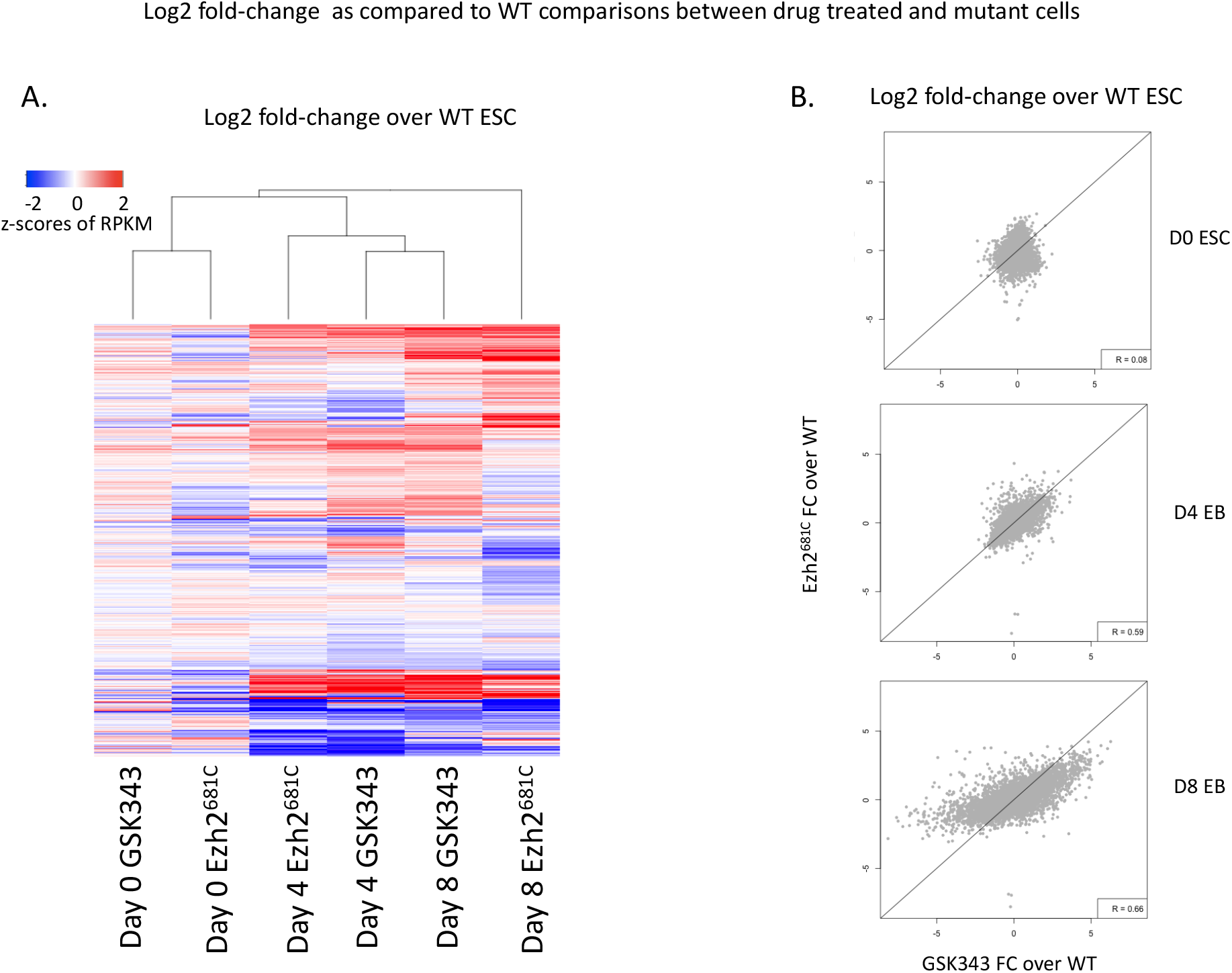
(A) Unsupervised clustering showing the fold change over WT ESC of GSK343 treated or Ezh2681C-99 cells. The day of differentiation is the stronger predictor of clustering. (B) Direct comparison of mutant and drug treated cells RNA-seq fold change over WT gene expression.

**Supp. Fig. 6.**
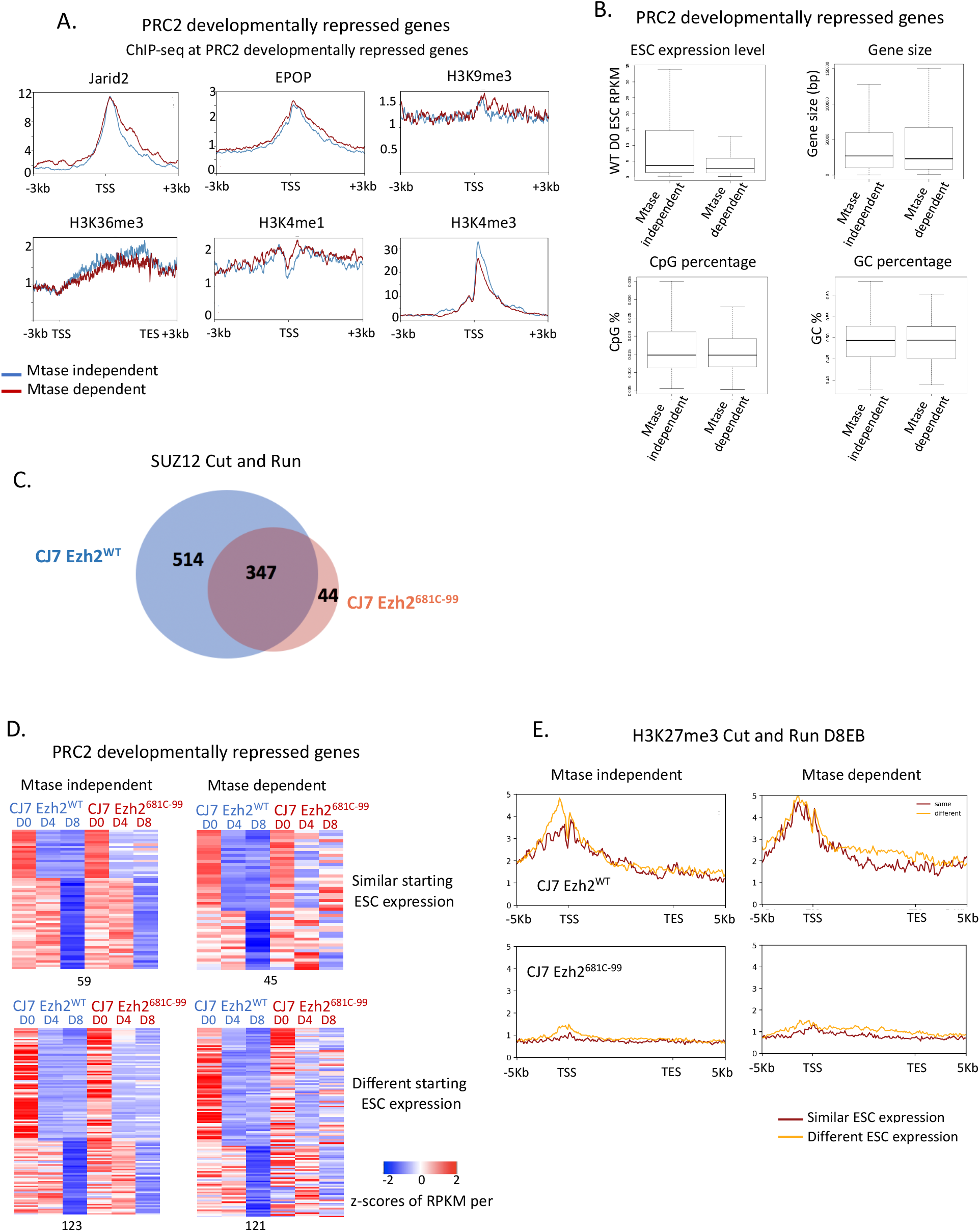
(A) Average profiles showing binding of other PRC2 components and histone marks in wild-type and mutant cells at Mtase dependent or independent and developmentally repressed PRC2 target genes over the time course. (B) Boxplots showing comparisons of different characteristics of Mtase dependent and independent, developmentally repressed PRC2 target genes in wild-type and mutant cells. (C) Venn diagram showing the overlap in SUZ12 peaks from CUT&RUN experiments from WT and Ezh2^618C-99^ cells. (D) Heatmaps showing developmentally repressed genes from Fig 3D, separated based on expression levels of wild-type and mutant ES cells at Day 0 of the time-course. Genes with similar expression levels at Day 0 in wild-type and mutant (within 1.2 fold) are on the top and those with different starting expression levels (more than 1.2 fold) are at the bottom. (E) Average profiles showing H3K27me3 binding in wild-type and mutant cells by cut-and-run at Mtase dependent or independent and developmentally repressed PRC2 target genes that have either different (more than 1.2 fold) or similar (within 1.2 fold) expression over the time course.

**Supp. Fig. 7.**
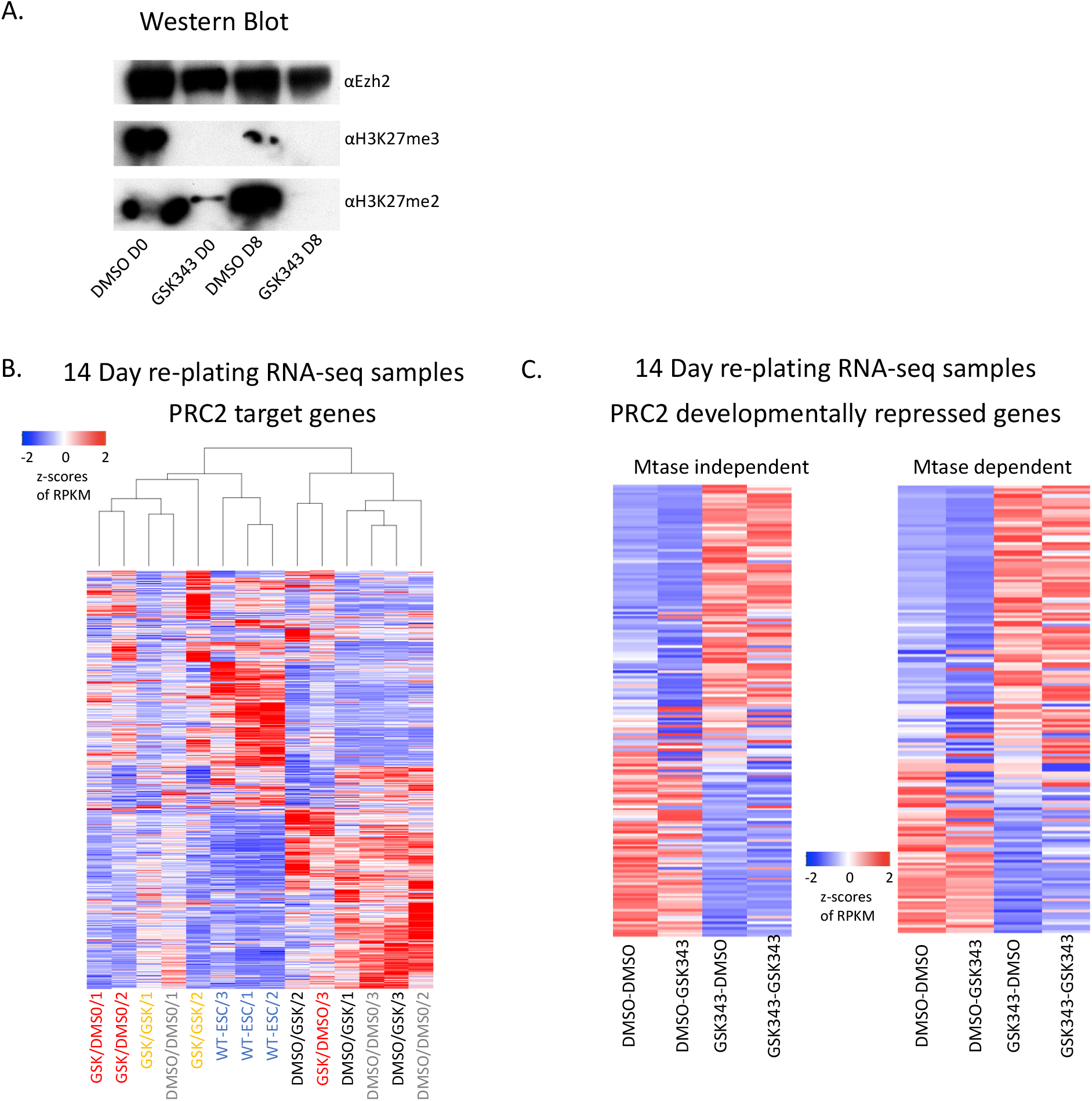
(A) Western blot of CJ7 cells treated with DMSO or GSK343 showing the (levels of Ezh2, H3K27me3 and H3K27me2 at Day 0 and Day 8 or EB formation. B) Unsupervised clustering of PRC2 target genes in re-plated cells and wild-type ES-cells. Untreated or drug treated cells were replated at 14 days in DMSO or GSK343. Z-scores of average RPKMs of triplicates (duplicates for GSK-GSK) are shown for each gene across all samples. (C) Heatmaps showing comparison of treated and untreated cells, re-plated in DMSO or GSK343 for previously identified Mtase dependent and independent genes. Z-scores of average RPKMs of triplicates (duplicates for GSK-GSK) are shown for each gene across all samples.

